# A Comparison of Methods to Measure Fitness in *Escherichia coli*

**DOI:** 10.1101/016121

**Authors:** Michael J Wiser, Richard E Lenski

## Abstract

In order to characterize the dynamics of adaptation, it is important to be able to quantify how a population’s mean fitness changes over time. Such measurements are especially important in experimental studies of evolution using microbes. The Long-Term Evolution Experiment (LTEE) with *Escherichia coli* provides one such system in which mean fitness has been measured by competing derived and ancestral populations. The traditional method used to measure fitness in the LTEE and many similar experiments, though, is subject to a potential limitation. As the relative fitness of the two competitors diverges, the measurement error increases because the less-fit population becomes increasingly small and cannot be enumerated as precisely. Here, we present and employ two alternatives to the traditional method. One is based on reducing the fitness differential between the competitors by using a common reference competitor from an intermediate generation that has intermediate fitness; the other alternative increases the initial population size of the less-fit, ancestral competitor. We performed a total of 480 competitions to compare the statistical properties of estimates obtained using these alternative methods with those obtained using the traditional method for samples taken over 50,000 generations from one of the LTEE populations. On balance, neither alternative method yielded measurements that were more precise than the traditional method.

## Introduction

The concept of fitness is central to evolutionary biology. Genotypes with higher fitness will tend to produce more offspring and thereby increase in frequency over time compared to their less-fit competitors. Fitness, however, is often difficult to measure, especially for long-lived organisms. Unlike traits such as color, fitness cannot be observed at a single point in time, but instead it must be measured and integrated across the lifespan of the individuals. Thus, researchers typically measure fitness components – such as the number of seeds produced or young fledged – and use them as proxies for fitness.

These limitations can be overcome in experimental evolution studies using microorganisms. Microbial systems typically have rapid generations and require little space, making them attractive for laboratory-based studies. Replicate populations founded from a common ancestor allow researchers to examine the repeatability of evolutionary changes. Environments can be controlled, reducing uninformative variation between samples or populations and allowing precise manipulations of conditions of interest. Also, one can often freeze microbial populations at multiple points along an evolutionary trajectory and revive them later, allowing direct comparisons between ancestral and derived populations [1,2]. Owing to these advantages, evolution experiments with microbes are becoming increasingly common [3–5]. Thus, it is important to be able to accurately quantify fitness in these experiments, in order to understand the evolutionary dynamics at work.

One commonly employed method of quantifying microbial fitness is to calculate the maximum growth rate (V_max_) of a culture growing on its own [6–10], usually by measuring the optical density of the culture over time. These measurements have the advantages of being simple and fast; a spectrophotometer can measure many samples in a multi-well plate in quick succession, and systems can be programmed to take measurements over the full growth cycle of a culture. However, maximum growth rate is typically only one component of fitness even in the simplest systems [11], and hence it provides, at best, only a proxy for fitness.

A second type of fitness measurement comes from studies where microbes are adapting to stressful compounds, such as antibiotics. In these situations, researchers typically quantify the Minimum Inhibitory Concentration (MIC) of the compound, and those organisms with higher MICs are considered to be more fit in environments that contain that compound, as it takes more of the substance to inhibit their growth [12,13].

A third approach for quantifying fitness in microbial systems—and the approach that most closely corresponds to the meaning of fitness in evolutionary theory—uses a competition assay. The basic approach is to compete one strain or population against another and directly measure their relative contributions to future generations. This approach typically produces a measure of relative, rather than absolute, fitness. Relative fitness is more important than absolute fitness when considering the evolutionary fate of a particular genotype, provided that absolute fitness is high enough to prevent extinction of the entire population [14,15]. Competitive fitness assays, by measuring the net growth of two different populations, incorporate and integrate differences across the full culture cycle, which may include such fitness components as lag times, exponential growth rates, and stationary phase dynamics in batch culture [11,16].

Despite their relevance to evolutionary theory, competitive fitness assays sometimes have practical limitations. In particular, and the focus of our paper, these measurements are more precise when the two competitors have similar fitness than when one is substantially more fit than the other. When one competitor is markedly less fit, its abundance will decrease over the course of the competition assay, potentially reaching values low enough that measurement error has a large impact. Thus, as the duration of an evolution experiment increases, and the fitness of the evolved organisms increases relative to the ancestral competitor, the measurement error also tends to increase, as we will show in this study.

We used a population from the Long-Term Evolution Experiment (LTEE) with *Escherichia coli* to investigate whether changes in the methods of performing competition assays – changes meant to reduce the discrepancy in the final abundances of the competitors – would yield more precise fitness measurements. The LTEE has been described in detail elsewhere [1,17–19], and a brief summary is provided in the Materials and Methods section below. Previous work in this system has established that changes in V_max_ explained much, but not all, of the improvement in relative fitness in this system, at least in the early generations [11].

## Materials and Methods

### Experimental conditions

The LTEE is an ongoing experiment that began in 1988, and which has now surpassed 50,000 bacterial generations. The experiment uses a Davis Minimal salts medium with 25 μg/mL glucose (DM25), which supports densities of ~3-5 x 10^7^ bacteria per mL. Each population lives in 10 mL of DM25 in a 50-mL glass Erlenmeyer flask incubated at 37C and shaken at 120 rpm. Every day, a member of the research team dilutes each population 1:100 into fresh media. This dilution sets the number of generations, as the regrowth up to the carrying capacity allows log_2_ 100 ≈ 6.64 cell divisions per day.

### Bacterial strains

The LTEE has 12 populations of *E. coli* [1]. Six populations were founded by a strain called REL606 [20] and six by the strain REL607. REL606 is unable to grow on the sugar arabinose (Ara^−^); REL607 is an Ara^+^ mutant derived from REL606. The DM25 medium does not contain arabinose, and the arabinose-utilization marker is selectively neutral in the LTEE environment [20]. In this study, we use both ancestral strains as well as samples taken from one population, called Ara-1, at generations 500, 1000, 1500, 2000, 5000, 10,000, 15,000, 20,000, 25,000, 30,000, 35,000, 40,000, 45,000, and 50,000. We also use a strain called REL11351, which is an Ara^+^ mutant of a clone isolated from the 5000-generation sample of population Ara-1.

### Fitness measurements

We quantify fitness in this system as the ratio of the realized growth rates of two populations while they compete for resources in the same flask and under the same environmental conditions used in the LTEE. This calculation is identical to the ratio of the number of doublings achieved by the two competitors. In all cases, we compete samples from the Ara-1 population (including the ancestor REL606) against an Ara^+^ competitor (either REL607 or REL11351). We distinguish the two competitors on the basis of their arabinose-utilization phenotypes; Ara^-^ and Ara^+^ cells produce red and white colonies, respectively, on Tetrazolium Arabinose (TA) agar plates [1,21].

We employ three different methods for measuring fitness in this study. For all three methods, we begin by removing aliquots of the competitors from the vials in which they are stored at –80C into separate flasks containing Luria-Bertani (LB) broth. The cultures grow overnight at 37C and reach stationary phase. We then dilute each culture 100-fold into 0.86% saline solution and transfer 100 μL into a flask containing 9.9 mL of DM25. These cultures grow for 24 h under the same conditions as the LTEE, so that all competitors are acclimated to this environment. We then jointly inoculate 100 μL in total of the Ara-1 population sample and the Ara^+^ competitor into 9.9 mL of DM25. We immediately take an initial 100-μL sample of this mixture, dilute it in saline solution, and spread the cells onto a TA plate. The competition mixture is then incubated in the same conditions as the LTEE for 24 h, at which point we take a final 100-μL sample, dilute it, and spread the cells onto a TA plate. We count each competitor on the TA plates, and multiply the numbers by the appropriate dilution factor to determine their initial and final population sizes. We calculate fitness as

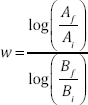

where *w* is fitness, *A* and *B* are the population sizes of the two competitors, and subscripts *i* and *f* indicate the initial and final time points in the assay.

For the Traditional method, we measure the relative fitness of the evolved population samples against the Ara^+^ ancestor, REL607. We inoculate the competition flasks with 50 μL (an equal volumetric ratio) of each competitor. This method has been used extensively in evaluating fitness in the LTEE [1,2].

The Altered Starting Ratio (ASR) method also uses the ancestral Ara^+^ strain as the common competitor. However, we inoculate the competition flasks with 20 μL of the evolved population and 80 μL of the ancestral population, leading to an initial 1:4 volumetric ratio. This difference in the starting ratio increases the population size of the ancestor at the end of the competition assay, which reduces the problem of small numbers when the ancestor is much less fit than the evolved population. The initial ratio is not so extreme, however, that it is difficult to enumerate the evolved population at the start of the competition assay. We attempted to keep total plate counts around a few hundred colonies, with at least 20 of the minority competitor, to reliably estimate population densities [22], and we chose this initial ratio with that objective in mind. It seemed particularly important to increase the final count of the ancestral population in the context of our fitness measurements; smaller numbers are subject to increased sampling error, and the realized growth rate of the ancestor is the denominator when calculating the relative fitness of the evolved population, which can magnify the measurement error.

Using the Different Common Competitor (DCC) method, we compete the evolved population samples against the marked clone from generation 5,000, rather than against the marked ancestor. We chose a 5,000-generation clone because its fitness was near the geometric mean of fitness values spanning generations 0 to 50,000, and thus might reduce the overall disparity in population counts across the full time series being considered. We inoculate the competitions with equal volumes (50 μL each) of the Ara-1 population sample and reference competitor. We considered that this method might increase the precision of our fitness measurements because the ratios used in the fitness calculation tend to be more precise as they approach 1.

We selected 15 time points from the focal population Ara-1 to evaluate these three methods: generations 0, 500, 1,000, 1,500, 2,000, 5,000, 10,000, 15,000, 20,000, 25,000, 30,000, 35,000, 40,000, 45,000, and 50,000. We ran competitions as complete blocks; each block included one competition for each time point using each method, plus an additional competition (see below) used as a scaling factor to compare the methods. We performed a total of 10 replicate blocks, and so there were a total of 450 competition assays to measure fitness (3 methods x 15 time points x 10 blocks) plus an additional 30 assays to generate the scaling factors.

A scaling factor was necessary for comparing the DCC method with the Traditional and ASR methods, because the DCC method measured fitness relative to a different competitor than the ancestor used for the other two methods. To calculate this scaling factor, we performed an additional competition between the Ara^−^ ancestor (REL606) and the Ara^+^ reference competitor (either REL607 or REL11351) for each method in every block. We then divided the fitness values from all of the competition assays for a given method and block by the fitness value that served as the scaling factor. We did not otherwise include the scaling-factor competitions in our data analysis. We applied the same procedure to all three methods to ensure consistency, although adjusting for the scaling factor was not otherwise required for the Traditional and ASR methods.

The data and analysis scripts are available at the Dryad Digital Depository (doi pending acceptance).

### Statistical methods

We performed statistical analyses in R version 2.14.1. We fit the fitness trajectories using nonlinear least-squares regression, as implemented with the nls() function. We performed ANOVAs using the aov() function. For the single-generation ANOVAs, Method was a fixed factor and Block was a random factor. For the combined ANOVA, Generation was included as a fixed factor.

### Bootstrapping

We employed a bootstrap procedure to compare the differences between the coefficients of variation in our three methods to a null distribution. We sampled the total dataset with replacement, to produce 3 datasets of equal size, each containing 10 measurements at each generation. We then fit a linear regression of the coefficient of variation against time (i.e., generation) to each of the 3 datasets. We then summed the squares of the differences between each pairwise combination of the 3 linear regressions over all 15 time points when fitness was measured. We repeated this entire procedure 1,000,000 times, and we compared the observed sum of the squared differences to this distribution.

## Results and Discussion

There are two fundamental ways in which these different methods could produce meaningfully different results. One way is that different methods could produce significantly different fitness estimates. In that case, we would need additional information or another criterion to determine which method was superior. The other way is that different methods could have different levels of precision; that is, one method may have significantly less variation in measured values across replicate assays than another. In this case, the method with the greater precision would clearly be preferred.

Fig. 1 shows the results of our fitness assays for all three methods, with trajectories fit to the data obtained using each method. These trajectories are in the form of an Offset Power Law:

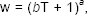

where w is fitness, T is time in generations, and *a* and *b* are model parameters, as derived in [2]. All three methods produce virtually identical fitness trajectories. S1 Table shows the results of ANOVAs performed at each generation to test for variation among the three methods in the mean fitness values they produce; the effect of Method was not significant in any of the 15 tests, even without accounting for multiple tests. From these results, we conclude that the three methods do not produce meaningfully different estimates of mean fitness.

**Figure 1:**
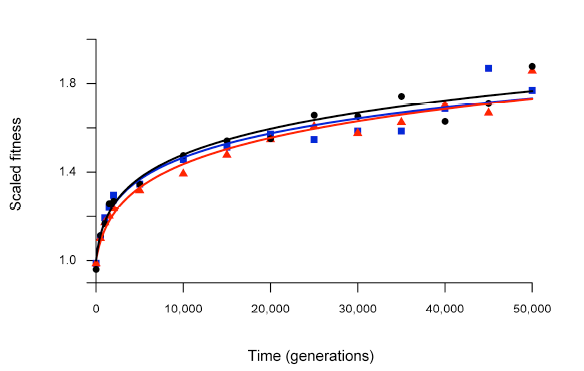
Fitness trajectories over time. Fitness trajectories for each method, shown separately, have the form w = (*bT* +1)^*a*^, where w is fitness, T is time in generations, and *a* and *b* are model parameters. Black circles and curve show the Traditional method; blue squares and curve show the ASR method; red triangles and curve show the DCC method.

Next, we calculated the coefficient of variation (i.e., the standard deviation divided by the mean) for each method at each time point to determine whether they differed in their precision. We then constructed a linear model of the coefficient of variation as a response to time (i.e., generation) and method. Fig. 2 shows the data and linear model fit to the coefficients of variation for all three methods. Table 1 presents the ANOVA table for this model. There is a highly significant tendency for the coefficient of variation to increase in later generations, as the evolving bacteria become progressively more fit, as discussed in the Introduction. However, the effect of Method was not significant as a predictor of the coefficient of variation, although a p-value of 0.0762 is suggestive. On inspection of the data (Fig. 2), it is clear that any difference between the methods is driven by the ASR method having a higher coefficient of variation – and thus lower precision – in early generations. Consistent with that appearance, when we removed the ASR method from the analysis and performed an ANOVA on the remaining data, there was no suggestion of any difference between the Traditional and DCC methods (Table 2, p = 0.8802).

**Figure 2:**
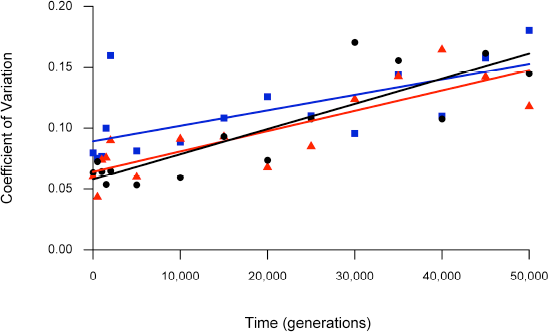
Coefficient of variation over time. Lines are linear regressions on the relevant data. Black circles and line show the Traditional method; blue squares and line show the ASR method; red triangles and line show the DCC method. S1 Fig. shows the confidence bands associated with each regression line.

**Table 1:** ANOVA on the coefficient of variation across time and comparing the three methods used to estimate fitness.

|  | df | SS | MS | F | p |
| --- | --- | --- | --- | --- | --- |
| Time | 1 | 0.03672 | 0.03672 | 69.664 | <0.0001 |
| Method | 2 | 0.00289 | 0.00145 | 2.743 | 0.0762 |
| Residuals | 41 | 0.21610 | 0.00053 |  |  |

**Table 2:** ANOVA on the coefficient of variation across time and comparing the Traditional and DCC methods.

|  | df | SS | MS | F | p |
| --- | --- | --- | --- | --- | --- |
| Time | 1 | 0.03068 | 0.03068 | 70.035 | <0.0001 |
| Method | 1 | 0.00001 | 0.00001 | 0.023 | 0.8802 |
| Residuals | 27 | 0.01183 | 0.00044 |  |  |

We can also express the differences between these methods as follows. The regression line for the coefficient of variation based on the ASR method is always higher than at least one of the other two methods (Fig. 2), and therefore it is never the best method, at least for the system and generations analyzed here. By contrast, the Traditional and DCC methods yield coefficients of variation, as inferred from the regression lines, that are very similar and always within the 95% confidence interval of one another (S1 Fig). Which of these two methods gave a lower point estimate of the coefficient of variation varied over time, but the difference was not significant (Table 2).

An alternative way to assess whether the differences in the coefficient of variation between the methods are statistically significant involves bootstrapping the data, as detailed in the Methods section. Fig. 3 shows that the observed differences in the coefficient of variation among the three methods are no greater than would be expected by chance if there were no differences among the methods.

**Figure 3:**
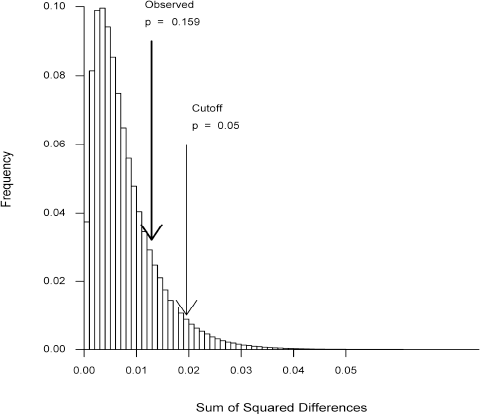
Histogram of bootstrap analysis. Histogram showing the distribution for the bootstrapped sums of squared differences in the coefficient of variation for 3 arbitrary groupings of the combined data. The dark arrow indicates the difference for the actual grouping of the 3 methods employed. The light arrow shows the most extreme 5% of the sums of the squared differences.

Over the range of fitness changes that we observed in the LTEE, neither alternative method for assaying fitness (ASR or DCC) outperformed the Traditional method. Given its extensive prior use in this study system [1,2,17], we therefore prefer to use the Traditional method for fitness competitions that span this range. It is important to note, however, that the ASR or the DCC method might turn out to have higher precision in systems that exhibit larger fitness changes than the system studied here, as suggested by the regression lines in Fig. 2. The LTEE has, to our knowledge, run for many more generations than any other evolution experiment, but the extent of fitness improvements has been less than that seen in some other shorter-duration experiments. The relatively limited fitness gains that have occurred during the LTEE reflect the fact that the experimental environment is quite benign; also, the ancestor of the LTEE had been studied by microbiologists for many decade [23] and was thus probably already well-adapted to general laboratory conditions. Other experiments conducted for fewer generations, but performed under more stressful conditions or founded by less-fit ancestors, might reach fitness differences where these or other alternative methods would be helpful. Table 3 summarizes the duration and range of fitness improvements reported in a number of other evolution experiments that used a variety of microorganisms including bacteria, fungi, and viruses (see also Table 2.3 in [24]).

**Table 3:** Selected evolution experiments

| Reference | Organism | Generations | Wf / Wi | Wf - Wi |
| --- | --- | --- | --- | --- |
| This study | *E. coli* | 50,000 | 1.88 | 3.5 / day |
| [25] | *E. coli* at 32C | 2,000 | 1.10 |  |
|  | *E. coli* at 42C | 2,000 | 1.19 |  |
| [26]* | *E. coli* | 1,100 | 1.98 | 0.23 / h |
| [27]** | *Saccharomyces cerevisiae* | 300 | 1.80 |  |
| [28] | *Aspergillus nidulans* | 800 | 1.48 |  |
| [29] | phage Φ6 with bottleneck = 10 | 100 | 1.26 | |
| | phage Φ6 with bottleneck = 1,000 | 40 | 2.03 | |
| [30] | phage G4 | 180 | 1.18 | 3.8 / h |
|  | phage ID2 | 600 | 2.55 | 13.5 / h |

\* Mean calculated from four replicate populations
\*\* Value estimated from figure

## Conclusion

We performed 480 assays to compare three different methods for estimating the relative fitness of bacterial competitors. The three methods generated results that were not meaningfully or significantly different in terms of either their mean values or dispersion. The only suggestion of a meaningful difference was that the ASR method appeared worse than the other two methods in the early generations, when the fitness gains of the evolved bacteria were still fairly small. Therefore, we see no compelling reason to adopt one of the alternatives to the Traditional method when analyzing systems that have achieved fitness gains less than or similar to those measured in the LTEE over its first 50,000 generations.

## Acknowledgments

We thank C. Turner, A. Lark, R. Maddamsetti, and C. Strelioff for helpful discussions during manuscript preparation and N. Hajela for technical assistance. This work was supported by grants from the National Science Foundation (DEB-1019989) including the BEACON Center for the Study of Evolution in Action (DBI-0939454), and by the Hannah Chair Endowment at Michigan State University. The data and analysis scripts are available at the Dryad Digital Repository (doi available upon acceptance).

## Supporting Information Captions

S1 Figure: Temporal trends in the coefficient of variation across replicate assays for the three different methods used to measure fitness. Black circles show the Traditional method; blue squares show the ASR method; red triangles show the DCC method. The solid colored lines show the linear regressions based on the corresponding data. The dashed colored curves show the 95% confidence bands for the regressions for the three methods: A) Traditional, B) ASR, and C) DCC. The points and regression lines are the same across all three panels, but the confidence bands are shown separately for clarity.

S1 Table: ANOVAs of fitness for three methods, by generation. Analyses of variance of measured fitness values for the three methods, analyzed separately for the various generations examined.

